# Native ribonucleases process sgRNA transcripts to create catalytic Cas9/sgRNA complexes *in planta*

**DOI:** 10.1101/2020.02.06.937003

**Authors:** Will B. Cody, Herman B. Scholthof

## Abstract

The current CRISPR/Cas9 gene editing dogma for single guide RNAs (sgRNA) delivery is based on the premise that 5′ and 3′ nucleotide overhangs negate Cas9/sgRNA catalytic activity *in vivo*. This has led to engineering strategies designed to either avoid or remove extraneous nucleotides on the 5′ and 3′ termini. Previously, we used a *Tobacco mosaic virus* viral vector to express both GFP and a sgRNA from a single virus-derived mRNA in *Nicotiana benthamiana*. This vector yielded high levels of GFP and catalytically active sgRNAs. Here, in an effort to understand the biochemical interactions of this result, we used *in vitro* assays to demonstrate that nucleotide overhangs 5′, but not 3′, proximal to the sgRNA do in fact inactivate Cas9 catalytic activity at the specified target site. Next we showed that *in planta* sgRNAs bound to Cas9 are devoid of the expected 5′ overhangs transcribed by the virus. Furthermore, when a plant nuclear promoter was used for expression of the GFP-sgRNA fusion transcript it also produced indels when delivered with Cas9. These results reveal that 5′ “auto-processing” of progenitor sgRNAs occurs natively in plants. Towards a possible mechanism for the perceived “auto-processing”, we found, using *in vitro* generated RNAs and those isolated from plants, that the 5′ to 3′ exoribonuclease XRN1 can degrade elongated progenitor sgRNAs whereas the mature sgRNA end-products are resistant. Comparisons with other studies suggest that sgRNA “auto-processing” may be a phenomenon not unique to plants, but other eukaryotes as well.

**Summary:** Native *Nicotiana benthamiana* ribonucleases cleave exogenous nucleotides 5′ to the sgRNA spacer transcript to create catalytic Cas9/sgRNA complexes *in planta*.

## Introduction

The CRISPR/Cas9 platform, found natively in *Streptococcus pyogenes*, has been developed into a diverse set of functional genetic tools, used in gene editing technology (Mali et al. 2013; Cong et al. 2013) and transcriptional control through gene activation (Gilbert et al. 2013) or repression (Qi et al. 2013). All of these technologies rely on the single guide RNA (sgRNA) programmable endonuclease Cas9 for specificity. One central restriction on the deployment of these CRISPR tools in eukaryotic organisms is the ability to simultaneously deliver both the Cas9 nuclease and sgRNAs in living cells. To circumvent this problem, viral vectors have been used in a wide variety of organisms for delivery of the protein and RNA products in adequate concentrations for a phenotypic response (Baltes et al. 2014; Malina et al. 2013; Platt et al. 2014). However, the long standing assumption that sgRNA delivery requires specific engineering for removal of 5′ and 3′ nucleotide overhanging sequences present in original progenitor transcripts, that are unrelated to the functional sgRNA sequence, has somewhat handicapped the convenient application and perhaps the efficiency of the technology.

Previously we aimed to create a sgRNA delivery system using the *Tobacco mosaic virus* based vector, TRBO, where we examined different sgRNA delivery strategies designed to eliminate nucleotide overhangs 5′ proximal to the spacer sequence and 3′ proximal to sgRNA scaffolding through the use of auto-catalytic ribozymes (Cody et al. 2017). Contrary to typical CRISPR dogma, the sgRNA without ribozymes –therefore containing lengthy nucleotide overhangs upstream and downstream of the sgRNA sequence resulting from the viral subgenomic transcriptional promoter–performed optimally in comparison to the ribozyme containing versions. Following this observation, we challenged the TRBO system to co-deliver a protein coding region (GFP) and a sgRNA using a single viral transcript (subgenomic mRNA), which resulted in a high incidence of genomic indels, and GFP expression levels comparable to the only GFP coding TRBO construct. These results, in addition to results from another study (Mikami et al. 2017), and other serendipitous findings (Ali et al. 2015; Cong et al. 2013), contradict the consensus in the field which suggests that *in vivo* delivered sgRNAs containing nucleotide overhangs prevent either Cas9-sgRNA complex assembly or its catalytic activity.

The activity associated with *in planta* produced subgenomic RNAs from the TRBO vector carrying both 5’ and 3’ overhangs suggest the existence of one or more of the following three activities that could explain Cas9 mediated formation of double-stranded breaks (DSBs) in plants such as *Nicotiana benthamiana*: 1) Cas9 tolerates large non-complementary overhangs 5’ proximal to the spacer and 3’ to the scaffold RNA, 2) Cas9 has the ability to cleave sgRNA overhangs, or 3) endogenous ribonucleases cleave sgRNA overhangs *in vivo*. The latter being the most popular implication which has been the suggested method in bacterial systems (Deltcheva et al. 2011; Karvelis et al. 2013). However, sgRNA 5′ overhang processing has not been supported through the identification of an associated protein or Cas9-protein complex in the bacterial models harboring native CRISPR systems (Karvelis et al. 2013; Deltcheva et al. 2011; van der Oost et al. 2014). In eukaryotic organisms, a considerable amount of energy focused on the delivery of sgRNAs that enable certain 5′ processing capabilities (Xie et al. 2015; Cermak et al. 2017a). Due to TRBO being an efficient protein and sgRNA co-delivery tool in *N. benthamiana*, as well as the overall lack of knowledge of native 5′ CRISPR RNAs (crRNA) processing currently in the literature –synonymous to 5′ sgRNA processing used in our models here –we aimed to better understand what is occurring to the 5′ end of sgRNAs *in vivo*, specifically in the experimental model *N. benthamiana*.

We first use *in vitro* assays with Cas9 and sgRNA transcripts containing nucleotide overhangs (nucleotides not corresponding/aligning to the 100 nucleotide chimera sgRNA sequence) of either or both the 5′ and 3′ ends of the sgRNA. In doing so, we concluded that in accordance with the prevailing dogma, 5′ overhangs do indeed completely inhibit the ability of the Cas9-sgRNA complex *in vitro*. Following these results we hypothesized, and found, that upon co-infiltration of a Cas9 and the TRBO-GFP-sgRNA co-expression constructs in *N. benthamiana* that Cas9 bound transcripts were enriched for sgRNAs which lacked the originally fused 5′ transcript sequences, indicative of a 5′ RNA processing event. Further sub-cellular fractionation analysis determined that the removal/processing of the 5′ nucleotide overhangs occurred in the plant cytosolic fraction. Next, we generated GFP-sgRNA transcripts that mimicked those generated by the viral system, but used a nuclear promoter for transcript expression. The results demonstrated that these transcripts were also capable of programming a catalytically active Cas9-sgRNA complex. Finally, to understand a potential RNA degradation pathway responsible for processing sgRNAs, we used both *in vitro* and *in planta* transcribed sgRNA templates and subjected them to the 5′ to 3′ exonuclease, XRN1. This did result in degradation of elongated RNAs but not of maturated sgRNA specific templates. These experiments directed us to develop a tentative model for creating catalytically active Cas9/sgRNA complexes that we believe could be applicable to other eukaryotes, and possibly the native processing system in *S. pyogenes*. Ultimately, the results from these experiments may have far reaching impacts on the development of CRISPR technology in future applications, but it also serves as a model for understanding fundamental biology of the native CRISPR/Cas9 5′ processing system.

## Results

### Viral subgenomic RNAs containing 5’ overhangs sgRNAs transcripts negate catalytic activity of Cas9 *in vitro*

To test whether Cas9 can cleave a protospacer-harboring DNA template using a sgRNA containing overhangs on either the 5’, 3′, or both ends, we elected to use the viral-based protein and sgRNA overexpression tool we previously developed, TRBO-G-3’gGFP (**Fig. 1A**), as a template to conduct *in vitro* Cas9 cleavage assays (Cody et al. 2017). In addition to carrying 5′ and 3′ UTR regions, TRBO-G-3’gGFP contains both a GFP protein coding segment and sgRNA targeting the *mgfp5* gene (gGFP). To model the subgenomic RNAs being produced from TRBO- G-3’gGFP, a T7 promoter carrying a forward primer was designed at the native coat protein subgenomic RNA transcription start site (T7-F1) and at the start of the spacer sequence of gGFP (T7-F2) to replicate both 5′ overhang carrying sgRNA and “clean” sgRNA (lacking extraneous nucleotides), respectively (**Fig. 1B**). To evaluate 3′ overhang effects on Cas9 nuclease activity, reverse primers were designed both in the 3′ TMV-UTR and on the 3′ most end of the sgRNA scaffolding, to replicate both 3′ overhang carrying sgRNA and “clean” sgRNA, respectively (**Fig. 1B**). PCR amplification of 5’ overhang carrying (T7/F1-R2), 3′ (T7/F2-R1), 5′ and 3′ (T7/F1-R1), and “clean” gGFP (T7/F2-R2) were used as a template for T7 transcription reactions followed by loading into purified Cas9 protein. Genomic DNA from the *mgfp5* harboring transgenic *N. benthamiana* 16c plants served as a template for amplification of *mgfp5* and amplicons were subsequently used as the DNA target for *in vitro* assays. A successful Cas9 cleavage is merited by the presence of digested DNA template that only occurred, in this case, when using a gGFP transcript without 5′ overhangs (**Fig. 1C**). Surprisingly, Cas9 still cleaved target DNA with a long 3′ gGFP nucleotide overhang *in vitro* (**Fig. 1C**). These results indicate that while 3′ sgRNA overhangs can be present and still allow for Cas9 dependent DSBs, sgRNAs carrying 5′ spacer sequence-adjacent overhangs inhibit Cas9 DNA cleavage.

**Figure 1.**
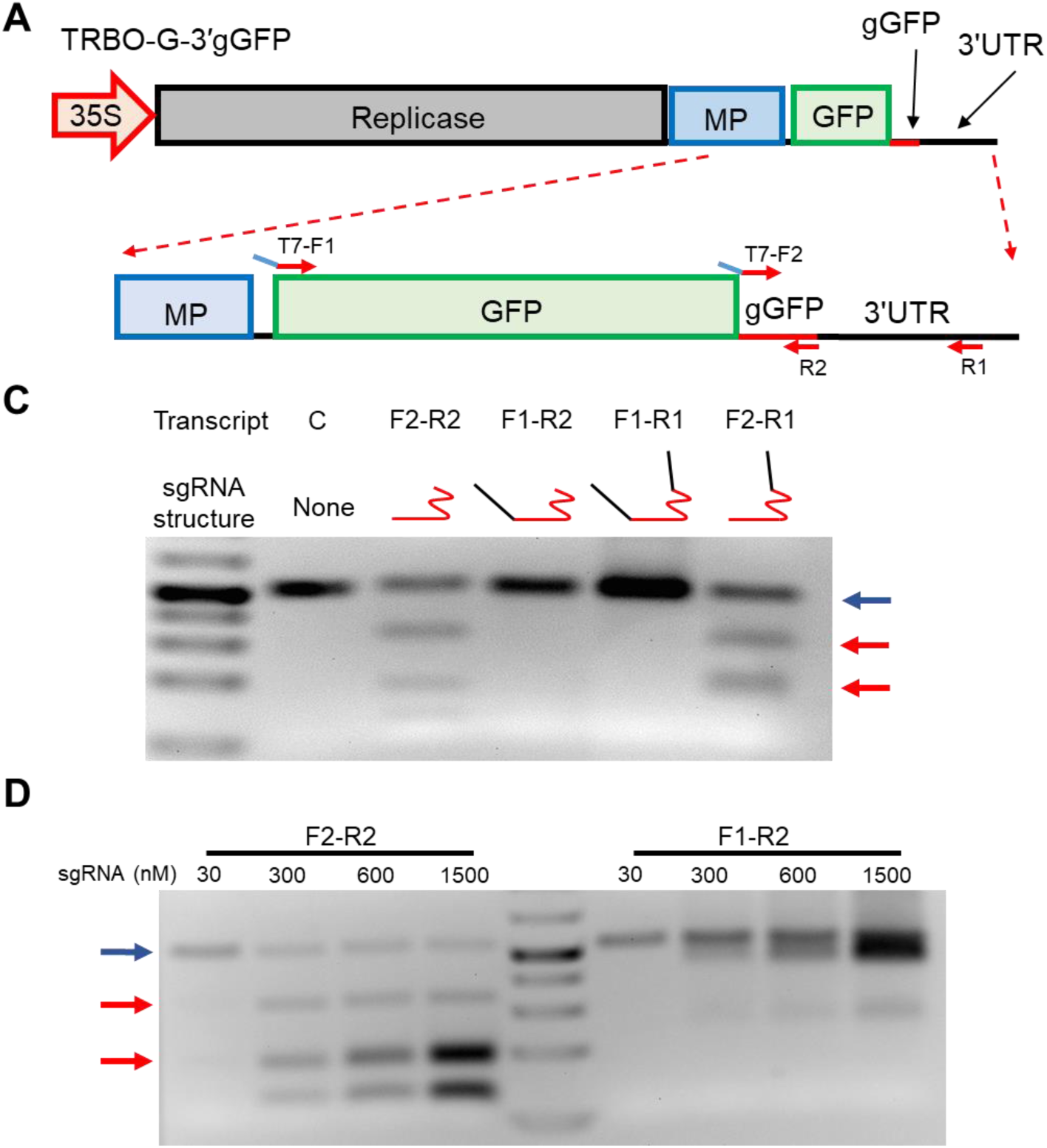
*In vitro* Cas9 cleavage assays using sgRNA with 5’ and 3’ nucleotide overhangs mimicking that of the TRBO subgenomic RNA. **A)** Genomic depiction of the TRBO protein and sgRNA delivery tool TRBO-G-3’gGFP. **B)** A zoomed in view of the TRBO-G-3’gGFP genome to illustrate T7 promoter carrying primers (blue lines with red arrows) used for T7 transcription reactions. Primers (red arrows) were designed to test if *in vitro* RNA products can direct Cas9 based DSBs. Blue lines upstream of the “T7” marked promoters represent the T7 promoter sequence used for transcription reactions. **C)** T7 transcription reactions using the corresponding primers as a template. Arrows indicate DNA fragments, with the blue arrow representing undigested *mgfp5* DNA template and red arrows represent cleaved *mgfp5* DNA template. Lanes are as follows: *mgfp5* DNA template and Cas9 nuclease only (C), gGFP without overhangs (F2- R2; red squiggle), gGFP with a 5’ overhang (F1-R2; black line), gGFP with both 5’ and 3’ overhangs (F1-R1), and gGFP with a 3’ overhang (F2-R1). **D)** sgRNA dosage depended cleavage of *mgfp5* DNA template using increased concentrations of gGFP without overhangs (F2-R2) and gGFP with a 5’ overhang (F1-R2). Bands that do not correspond to either undigested DNA or digested DNA (red and blue arrows) are sgRNA transcripts used in the assay.

Concentrations of sgRNAs in the previous Cas9 cleavage assays (**Fig. 1C**) were used at levels suited for optimal function of “clean” sgRNAs. To mimic the TRBO-sgRNA delivery system, which produces an abundance of sgRNAs *in planta*, and to rule out the possibility of sgRNA dosage-dependent Cas9 DNA catalysis events, we further examined *in vitro* catalytic activity using increasing concentrations (30 nM-1500 nM) of 5′ overhang carrying sgRNAs (T7/F1-R2) using the Cas9 *in vitro* cleavage assay system. The results demonstrated that even with large concentrations of 5′ overhang progenitor gGFP template available for Cas9 loading, there was no evidence of DNA cleavage (**Fig. 1D**). These results indicate that the increased concentrations of 5′-elongated-gGFP-progenitors observed with TRBO delivery *in planta* is unlikely the source of efficient Cas9 editing, but instead that native 5’ sgRNA processing abilities most likely exist *in planta*.

### Cas9 bound sgRNAs have processed 5′ ends *in planta*

Previously we established that co-delivery of pHcoCas9 (**Fig. 2A**) and TRBO-G-3’gGFP (**Fig. 1A**) results in the assembly of catalytically competent Cas9-sgRNA complexes *in planta* (Cody et al. 2017). However, *in vitro* results indicate that full length, TRBO generated, subgenomic RNA transcripts could not form catalytically active Cas9-sgRNA complexes. These results suggest that progenitor-sgRNA 5′ ends are being removed (processed) *in planta* by host factors, to produce sgRNAs end-products capable of targeted cleavage. To better understand the structure and composition of sgRNAs bound to Cas9 *in planta* and to determine if 5′ processing is occurring, immunoprecipitations of Cas9 from *N. benthamiana* 16c plants infiltrated with pHcoCas9, TRBO-G-3’gGFP, or pHcoCas9 and TRBO-G-3’gGFP were performed followed by RNA extractions. Additionally, an RT-PCR amplification scheme was designed using three primer sets to detect for an enrichment of TRBO-G-3’gGFP derived RNA product with a particular emphasis on shortened (*e.g*., processed) gGFP spacer fragments (**Fig. 2B**). Forward primers were designed in the genome of TMV as follows: i) within the movement protein (MP) coding segment (F1), ii) at the start codon of the downstream GFP (F2), and iii) at the 5′-end of the gGFP spacer sequence (F3). Since our earlier results demonstrated that 3′ sgRNA overhangs do not impede Cas9-sgRNA ability to induce DSBs (**Fig. 1C**), we elected to amplify sgRNA fragments using a reverse primer starting within the sgRNA scaffolding (R2) to enable us to focus on the biological relevant 5′ proximal to the spacer sequence.

**Figure 2.**
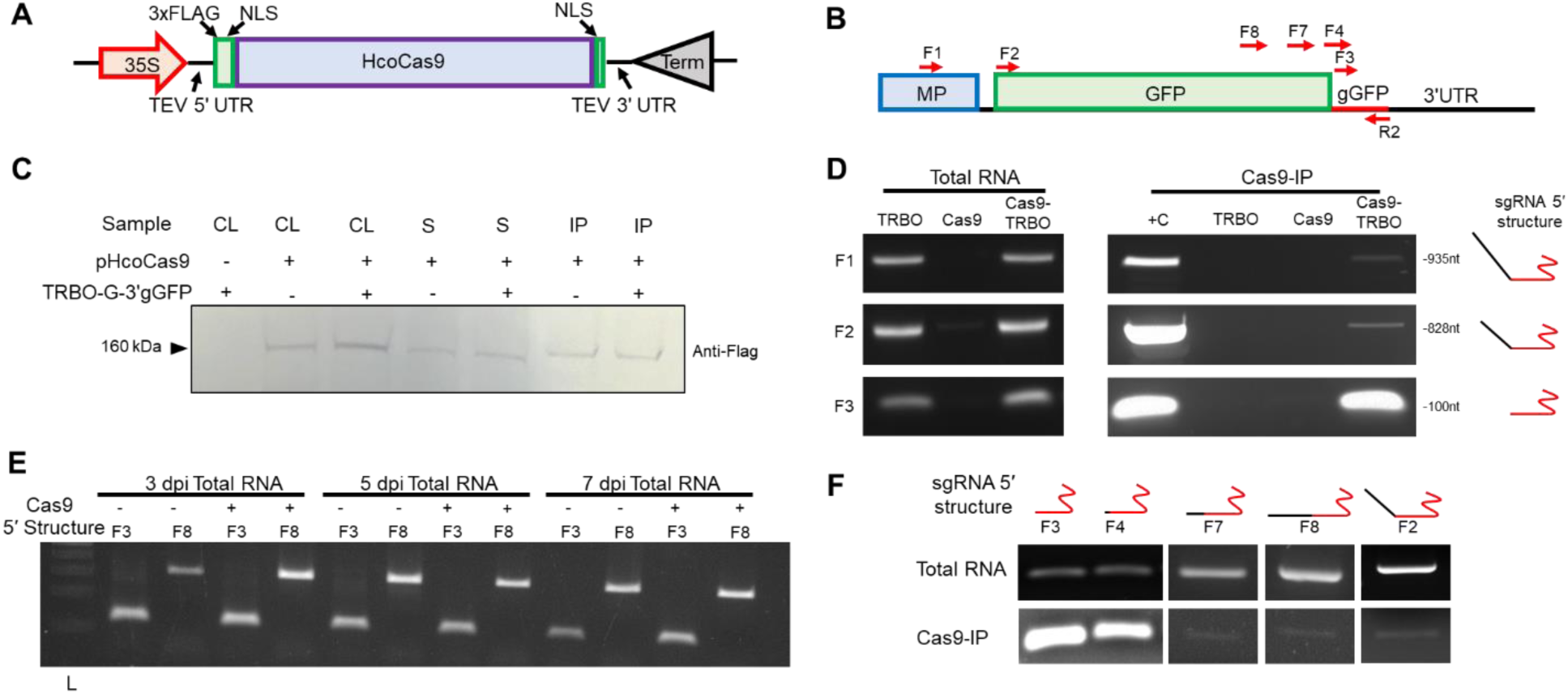
*In planta* sgRNA transcript processing, and Cas9 loading. **A)** The plant Cas9 expression construct pHcoCas9. This construct contains an N-terminal triple FLAG-tag (3xFLAG), nuclear localization signals (NLS), and a human codon-optimized Cas9 nuclease. Transcription is initiated by a CaMV 35S promoter and terminated by a 35S terminator. Transcripts also contain both a 5′ *Tobacco etch virus* (TEV) UTR and 3′ TEV UTR for increased translation efficiency. **B)** A zoomed in region of TRBO-G-3′gGFP depicting the 3′ genomic organization and corresponding primers (red arrows) used for RT-PCR experiments. Primers are designed to detect the 5′ condition of in vivo delivered gGFP products (presence of nucleotide overhangs or not). **C)** Western blot using anti-Flag antibody to detect Cas9 in cellular lysate (CL), Cas9-IP (IP), and supernatant from Cas9-IP before washing (S) upon infiltration with pHcoCas9 and/or TRBO-G- 3’gGFP, as indicated by + and -. **D)** RNA isolation was carried out using Cas9-IP samples for TRBO-G-3′gGFP (TRBO), pHcoCas9 (Cas9), and pHcoCas9/TRBO-G-3′gGFP (Cas9-TRBO). RT-PCR was performed using primers depicted in **A**, using cDNA from total RNA and Cas9-IP (also shown in **C**), to examine presence of in vivo gGFP 5′ overhangs. Enrichment of gGFP RNAs that do not encompass the predicted subgenomic RNAs as shown by ample amplification using F3-R2 primers and not F2-R2 for Cas9-TRBO. The positive control (+C) was carried out using TRBO-G-3′gGFP purified plasmid. The expected amplified sgRNA structure is indicated to the right with the red lines representing sgRNA specific sequence and black depicting viral RNA. **E)** Total RNA was sampled at 3, 5 and 7 dpi from 16c tissue infiltrated with TRBO-G-3′gGFP both with and without pHcoCas9. Total RNA was assayed for processed and unprocessed sgRNAs by using the F3 and F8 forward primer sets, respectively to detect different portions of the sgRNA containing TRBO transcripts. L indicates DNA ladder lane. **F)** RT-PCR using total RNA and Cas9 bound RNA (Cas9-IP) from 16c leaf tissue infiltrated with pHcoCas9 and TRBO-G-3′gGFP. Primers F4, F7, and F8 are located in increasing distance upstream to gGFP, respectively. Structures of 5′ sgRNA are depicted above the forward primer used with increasing length of black line representing longer 5′ overhang sequence.

Since we previously established that the majority of editing events occur during 2-3 days post-inoculation (dpi) (Cody et al. 2017), 3 dpi samples were assayed from each treatment for analysis. Cas9 protein was isolated through immunoprecipitation (IP) using a Cas9-specific antibody followed by protein G agarose bead pull-down. Cas9 protein isolation on the protein G agarose beads was verified via western blot detection (**Fig. 2C**). RNA extractions were carried out using all three Cas9-IP samples, and for comparison total RNA samples were also extracted for each tissue. RT reactions were performed using the sgRNA scaffold specific (R2) primer. Total RNA RT-PCR amplifications showed approximately equal quantities of product when comparing theTRBO-G-3′gGFP alone, versus the pHcoCas9 plus TRBO-G-3′gGFP co-infiltrated samples (**Fig. 2D**). Roughly equal expression quantities held true over 3, 5, and 7 dpi (**Fig. 2E**). In contrast, RT-PCR amplifications on the IP-products showed a clear enrichment of gGFP specific amplicons (F3-R2) in the pHcoCas9 and TRBO-G-3’gGFP co-infiltrated tissue compared to the predicted longer viral subgenomic RNA product (F2-R2) and genomic/first subgenomic containing RNA product (F1-R2) (**Fig. 2D**). In line with expectations, the two controls either devoid of sgRNA (pHcoCas9 alone) or Cas9 (TRBO-G-3’gGFP alone), did not yield amplification products of the expected molecular weight for each primer set.

We next aimed at testing whether processing specificity is manifested for sgRNAs loaded within Cas9 by examining if sgRNAs were specifically cleaved at the 5′ terminus of the mature sgRNA or if several subpopulations of sgRNAs containing various 5′ overhang lengths associated with Cas9. Towards this, forward primers were designed from the gGFP (F3) spacer sequence progressively moving upstream of the subgenomic RNA in increments (**Fig. 2B**). RT-PCR indicated a clear reduction in band intensity with primers used upstream and 5′ proximal to the gGFP spacer sequence (**Fig. 2F**). These data confirm that the 5′ end of gGFP is being processed (cleaved) *in planta* to eliminate the nucleotide overhang produced during viral subgenomic RNA production (transcription) with some level of specificity to the start of the 5′ spacer sequence. Furthermore, it appears that either Cas9 preferentially binds processed sgRNAs, or proper 5′ nucleotide removal is stimulated by association of Cas9 with the progenitor-sgRNA.

### TRBO synthesized sgRNAs bound to Cas9 are processed in the cytoplasm

Due to the finding that the majority of sgRNAs bound by Cas9 are processed, we next aimed at gaining an understanding of what host processes might be responsible for 5′ sgRNA maturation events *in planta*. One hypothesis based on other findings (Ohle et al. 2016; Sternberg et al. 2014) was that the mechanism of Cas9-sgRNA host DNA “target” scanning, demonstrated *in vitro*, involves the initiation of R-loop structures which can be processed by host RNase H enzymes. The manner in which Cas9 interrogates DNA to identify the protospacer sequence primarily relies on the 10 nt 3′ spacer “seed” sequence (Sternberg et al. 2014). When using TRBO-G-3′gGFP as the sgRNA delivery tool, in this system, there would be large RNA overhangs located upstream to the 5′ most nucleotide of protospacer complementary region, creating an R-loop structure which could potentially be recognized by RNase H enzymes. In other words RNase H-mediated processing of the progenitor-sgRNA would only occur in presence of a 100% complementary genomic protospacer sequence. To test this, we co-infiltrated pHcoCas9 and TRBO-G-3′gGFP into 16c *N. benthamiana* (containing *mgfp5*) plants and compared results with those obtained in wild-type (wt) *N. benthamiana* plants (not containing the genomic target). Following infiltration, 16c and wt plants were harvested at 3 dpi. Cas9 was immunoprecipitated (IP) and RNA was sampled from both wt and 16c Cas9-IP and the total lysate used for the IP reactions (**Sup Fig 1A**). RT-PCR products for total RNA lysate of 16c and wt plants indicated no discrepancies in band intensities using the previously designed primer sets (**Sup Fig 1B**). Following these results it was concluded that sgRNA 5′ processing was not reliant on a protospacer being present in the nuclear DNA and must be occurring through another mechanism.

To test if nuclear localization is required for sgRNA processing we removed the nuclear localization signals (NLS) from Cas9 and constructed p-NLSCas9 (**Sup Fig 1C**). 16c plants were then infiltrated with TRBO-G-3′gGFP as well as co-infiltrated with either the NLS lacking p- NLSCas9 construct or the NLS containing pHcoCas9 vector. To confirm a lack of localization to the genomic DNA, a proxy for nuclear localization, of the p-NLSCas9 encoded protein, 7 dpi DNA was assayed for verification of indel formation following each treatment. As expected, the pHcoCas9 construct produced DSBs from 16c genomic DNA whereas the p-NLSCas9 indel quantification resulted in levels undifferentiated from the TRBO-G-3′gGFP only control (**Sup Fig. 1D**). These results demonstrate that a lack of nuclear subcellular localization of Cas9 (– NLSCas9) negates complex catalysis of substrate DNA. Following these results, tissue was sampled from 4 dpi 16c plants and used for Cas9-IPs followed by RNA extractions as well as for total lysate RNA extractions. Total RNA and Cas9 bound RNA from both pHcoCas9 and p- NLSCas9 treatments were subject to RT-PCR and it was confirmed that sgRNA 5′ processing occurred in extracts containing either the NLS lacking or the NLS containing construct (**Sup Fig. 1E**).

These results indicated that DNA target recognition events in the nucleus might not be critical for progenitor-sgRNA processing, which suggests the possible contribution of cytoplasmic events to enable catalytic activity of the complex. Therefore, we next interrogated both the cellular localization of Cas9 protein and sgRNAs to identify the location of 5′ sgRNA processing (nucleus or cytosol). Using sub-cellular fractionation in combination with the previously developed RT-PCR scheme (**Fig. 2B**), we compared the fractions for relative levels of unprocessed full-length subgenomic RNAs to 5′ processed gGFP both with and without cellular production of Cas9 protein. For this, equal fractions were first analyzed for the presence of Cas9 protein through western blotting, which indicated that even though a sub-population of Cas9 protein accumulates in the cytosol Cas9 preferentially localizes to the nucleus (**Fig. 3A**). Both nuclear and cytosol fractions from pHcoCas9 and TRBO-G-3’gGFP co-infiltrated tissue were then used for Cas9-IP (**Fig. 3A**), followed by RNA extractions. Total RNA was also extracted from pHcoCas9 and TRBO-G-3’gGFP total cellular lysate as well as from the cytosol and nuclear lysate fractions. RT-PCR analysis from the total nuclear lysate and the Cas9-IP isolated from the nuclear fraction indicated that sgRNAs were being processed prior to translocation into the nucleus (**Fig. 3B**). Additionally, there was a clear enrichment for specific gGFP 5′ processed forms in the Cas9-IP cytosolic fraction reactions as compared to the reactions from the total RNA in cytosolic fraction (**Fig. 3B**). While cytosolic lysate showed no discrepancies between the gGFP processing forms, as in the total lysate control, the Cas9-IP RNA contained mostly 5′ processed forms of sgRNAs. Taken together, these data reinforce that 5’ sgRNA processing does not depend on Cas9 nuclear localization but instead occurs, at least primarily, in the cytosol using our viral-sgRNA delivery system.

**Figure 3.**
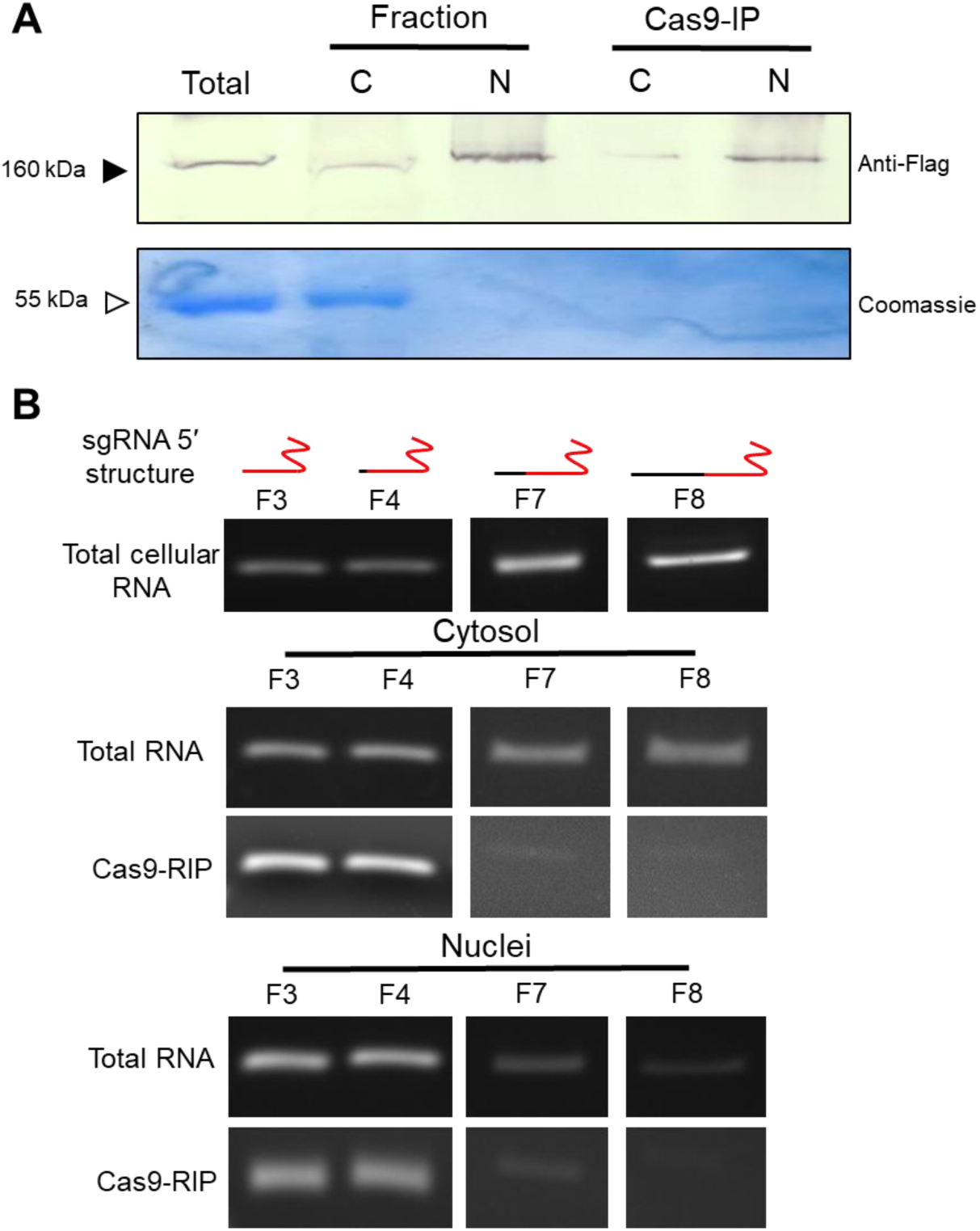
5′ processing of sgRNA transcripts of pHcoCas9 and TRBO-G-3′gGFP co-infiltrated 16c plants upon examination of total RNA and Cas9-IP RNA in nuclear and cytosolic fractions. **A)** Protein lysate and Cas9 immunoprecipitations (Cas9-IP) from total lysate (Total), cytosolic fraction (C) and nuclear fraction (N) were used for detection of Cas9. The top panel is a western blot to detect Cas9 (solid arrow 160 kDa) using an Anti-Flag primary antibody. The bottom panel is a Coomassie stain to detect Rubisco (open arrow 55 kDa). **B)** Samples assayed for Cas9 expression in **A** were used for detection of 5′ gGFP processing through RNA extractions followed by RT-PCR using forward and reverse primer sets described in **Figure 2A**. Total lysate RNA, total cytosolic RNA, Cas9-IP cytosolic RNA, total nuclear RNA, and Cas9-IP nuclear RNA were used to detect the cellular location of 5′ sgRNA processing. The expected sgRNA 5′ structure of the forward primers is depicted above the gel panels with the red structure representing the sgRNA and black lines depicting the size of the 5′ overhang amplified. Primer sets are as described in **Figure 2B**.

### Nuclear transcribed protein-sgRNA fusion transcripts create catalytically active Cas9 complexes

Next we aimed to understand if the 5′ sgRNA processing events are in some way associated with a host response to virus infection, or due to TMV (TRBO) replication and gene expression being localized to the cytosol. If cytosolic localization or virus infection response is in fact responsible for sgRNA 5′ processing events, then nuclear transcribed transcripts carrying nucleotide overhangs should not create catalytically active Cas9-sgRNA complexes. To examine this, we used the GFP-gGFP fusion transcript (progenitor-sgRNA), used in TRBO-G-3′gGFP, as a template and initiated its transcription from the *Arabidopsis thaliana* Pol III U6 nuclear promoter. The U6 promoter-based expression of sgRNAs retains transcripts to the nucleus, notably this quality was the reason the initial *in vivo* CRISPR/Cas9 assays used U6 promoter to drive sgRNA expression. However, in this case we used the nuclear localization of transcripts produced from the U6 promoter to discern the effect overhangs and specifically nuclear sgRNA overhangs have on successful host non-homologous end joining (NHEJ) double stranded break repair from a catalytically competent Cas9-sgRNA complex, as reflected by indel percentages.

To separate both cytosolic transcript expression/localization and potential viral host responses, the protein-sgRNA fusion transcript, U6-GFP-gGFP, was constructed along with a transcript producing only “clean” (no 5′ nucleotide overhangs) sgRNA, U6-gGFP, to serve as a control for DSB activity. Both U6-GFP-gGFP and U6-gGFP were inserted into the pHcoCas9 expression vector to produce pHco-U6-GFP-gGFP and pHco-U6-gGFP, respectively (**Fig. 4A**). Then, 16c plants were used for half-leaf assays using pHco-U6-GFP-gGFP and pHco-U6-gGFP to test for *in planta* catalytic activity (**Fig 4B**). Tissue was taken at 7 dpi from three assayed plant samples and subjected to PCR amplification followed by a *Bsg*I digestion. The pHco-U6-GFP- gGFP infiltrated tissue surprisingly showed substantial quantity of indels 17%-30%, but pHco- U6-gGFP was considerably higher at 33%-40% (**Fig. 4C**). Each half-leaf assay, indicated by number (**Fig. 4C**), consistently measured lower percentages of indel mutations using pHco-U6- GFP-gGFP compared to the pHco-U6-gGFP infiltrated part of the leaf (**Fig. 4D**). One possible explanation for the lower indel percentages using the pHco-U6-GFP-gGFP construct would be the length of the transcript (∼850 nts) being much longer than a typical Pol III transcribed RNA (100-150 nts), causing a decrease in gGFP expression due to lower levels of Pol III fidelity at the 3′ end of the transcript. To test if the discrepancy of indel mutation percentages between these two constructs was due to lower expression levels of the pHco-U6-GFP-gGFP transcripts or due to 5′ sgRNA overhangs impairing catalytic activity, 5 dpi half-leaf assays were used for RT-PCR expression analysis (**Fig. 4E**). Ultimately there was no difference in expression levels of gGFP between either pHco-U6-GFP-gGFP or pHco-U6-gGFP, indicating that the lower indel percentages from the pHco-U6-GFP-gGFP is due to a reduction in 5′ processing efficiency in host cells more than likely due to the extended nuclear localization of transcripts synthesized from U6 promoters, confirming that cytoplasmic localization stimulates progenitor-gRNA processing. Perhaps even more importantly these assays demonstrate that, in fact, pHco-U6- GFP-gGFP is capable of delivering sgRNAs with considerable 5′ overhangs that are clearly capable of producing indels in the presence of Cas9 and, as such, must have been processed to mature sgRNAs. Altogether, the results show that 5’ processing of progenitor sgRNA is not specific for viral-delivery but reflects a general event, thus contradicting the assumptions currently made in the literature on sgRNA function being incapacitated by substantial sgRNA nucleotide overhangs.

**Figure 4.**
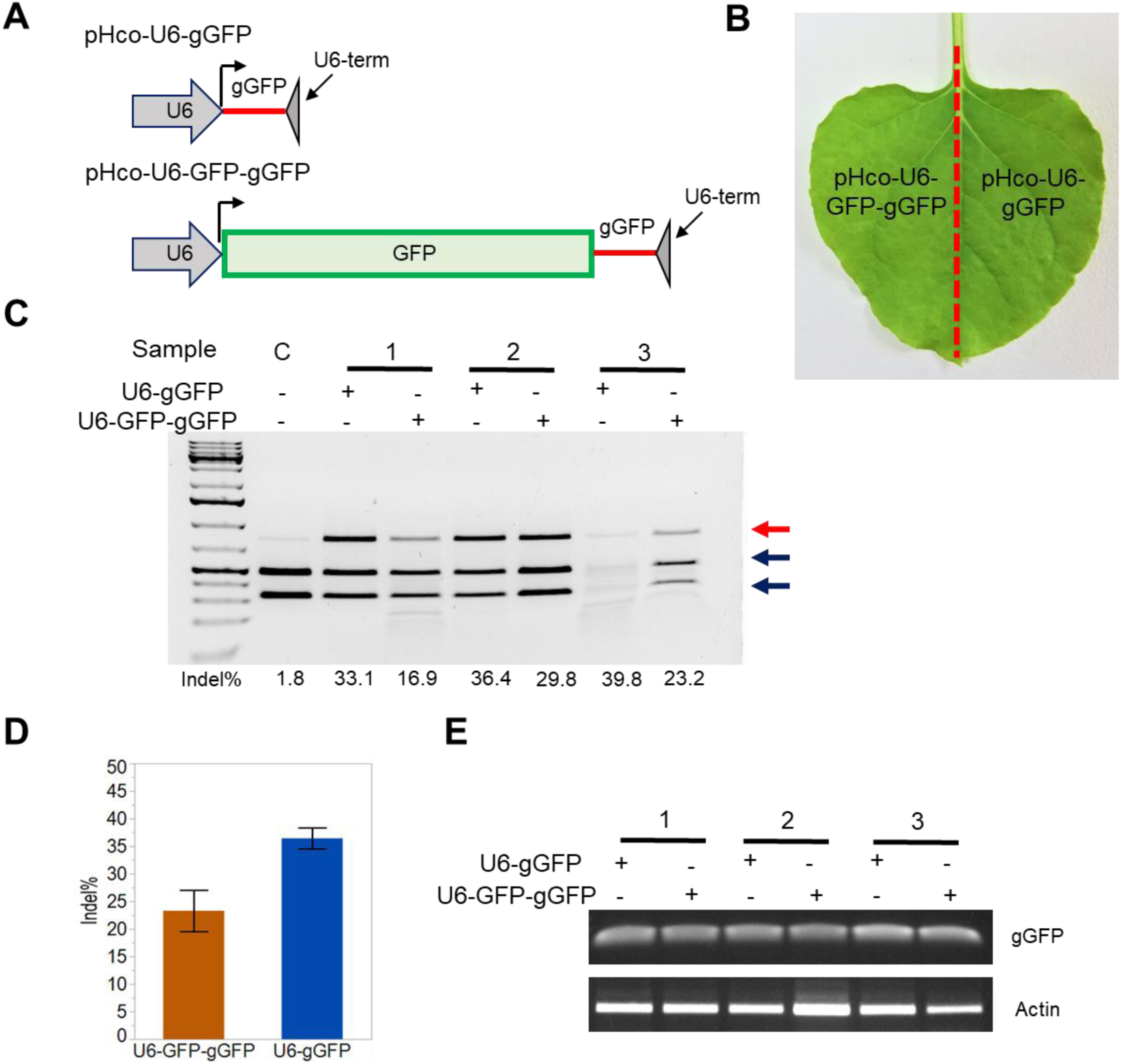
U6 nuclear promoter transcription of a sgRNA with a 5′ overhang *in planta*. **A)** Two sgRNA *in planta* delivery constructs cloned into pHcoCas9. pHco-U6-gGFP contains a polymerase III U6 promoter followed by the gGFP sgRNA and a U6 transcription terminator. pHco-U6-GFP-gGFP construct is same as pHco-U6-gGFP but contains the GFP protein coding region directly 5′ to the gGFP sequence and 3′ to the U6 promoter. **B)** Depiction of the experimental set-up of half-leaf assays used. Constructs shown in **A** were agroinfiltrated into one side of the leaf with the leafs midrib (red dashed line) serving to separate the treatments. **C)** *Bsg*I digest from *mgfp5* amplified PCR products of three replicates (Sample 1-3) half-leaf assays depicted in **B** and sampled at 7 dpi. The negative control (C) represents pHcoCas9 infiltrated 16c plants. The red arrow indicates *Bsg*I digest resistant bands containing indels and the blue arrows indicate digested, or wt *mgfp5* sequences. Indel percentages quantified using ImageJ image analysis software for each treatment are shown under the corresponding lane. **D)** Indel percentages calculated from **C** were used for one-way ANOVA statistical analysis. Mean indel value are represented by the colored bars for pHco-U6-GFP-gGFP (orange) and pHco-U6-gGFP (blue). Error bars represent the standard error of the mean (N=3; *P* = .035). **E)** RT-PCR analysis of half-leaf assays depicted in **B** used to compare the expression levels of gGFP from both the pHco-U6- gGFP and pHco-U6-GFP-gGFP. RNA was extracted at 7 dpi from the same leaves used in **C**. The top panel, gGFP, primers specifically amplifying gGFP expression were used. The bottom panel, Actin, primers specifically amplifying *N. benthamiana* Actin expression that was used as the loading control.

### *In vitro* and *in planta* generated sgRNAs are resistant to 5′ to 3′ exonuclease XRN1

While cytosolic expression of sgRNA transcripts carrying 5′ overhangs appears to be optimal for processing of gGFP into its final catalytically active form, we remained perplexed about how and what pathways might be involved. Due to the orientation of the RNA overhang on the sgRNA being 5′ proximal, we contemplated the possibility of a 5′ to 3′ exoribonuclease interacting with the sgRNA. The primary protein family responsible for this activity in eukaryotes is the XRN class of proteins. Indeed these proteins have also been characterized in other models to localize in both nuclear and cytosolic fractions, potentially explaining results seen in **Figure 4C** in regards to possible nuclear sgRNA processing activity. However, complications for experimenting with the XRN family of proteins in *N. benthamiana* are many due to the diversity of predicted *xrn* gene loci (**Sup Fig 2A**), and previous demonstration of their functional overlap in *Arabidopsis* (Kurihara 2017). However, one condition for typical XRN-family activity is the presence of a 5′ monophosphate (Jinek et al. 2011). Considering our TRBO subgenomic RNA generated sgRNA transcripts contain a 5′ cap structure, we speculated that cap removal might be a more reasonable genetic target due to the presence of only a single catalytic component, DCP2, in the native decapping complex and the high similarity of the two *dcp2* genes in *N. benthamiana* (Niben101Scf01105g01020.1 and Niben101Scf26315g00002.1)(Xu et al. 2006; Forment et al. 2005). However, our *Tobacco rattle virus* (TRV) viral induced gene silencing vector (VIGS) targeting *dcp2* transcripts (**Sup Fig 2B**) in fact increased DSBs at the target loci (**Sup Fig 2C**). This could be related to the a report that silencing of decapping enzymes, in fact, causes increased viral replication (Ma et al. 2015), and therefore these results could be a result of increased cellular content of sgRNA and Cas9.

Instead of moving forward with the rather complicated genetics of our *in planta* experimental model (**Sup Fig 2**) we looked towards recapitulating the XRN-sgRNA interaction *in vitro*. It was previously reported that tracrRNAs, coinciding with the 80 3′ most nucleotides on the sgRNA, are needed for the maturation (3′ and 5′ processing events) of crRNAs into catalytically competent crRNA/tracrRNA duplex (Deltcheva et al. 2011). Intriguing observations were that crRNAs are readily degraded in the native host *S. pyogenes* devoid of tracrRNAs, and tracrRNAs remain present in their active form in a strain without crRNAs. Perhaps this is due to an inherent stability of the tracrRNAs molecules? Indeed when exogenous TEX (5′ to 3′ exonuclease) is supplied to RNA from *S. pyogenes* lysate containing the predicted crRNA binding form of tracrRNA remains present while crRNAs are readily digested, indicating resistance to the enzyme (Deltcheva et al. 2011). Based on this, we hypothesized that in our *in planta* system, in contrast to the susceptible progenitor sgRNAs, the mature sgRNAs exhibit resistance to 5′ to 3′ processing enzymes such as XRN-1 (yeast) *in vitro*, towards an explanation for the processing phenomenon we see *in planta*.

In vitro assays were set up by supplying an exogenous RNA 5′ pyrophosphohydrolase (RppH) to produce a 5′ monophosphate transcript that can then readily be degraded by XRN proteins, in this case XRN-1. To test our hypothesis we generated two transcripts, one from the full predicted subgenomic RNA produced from TRBO-G-3′gGFP (F1-R2) and the other containing only the sgRNA sequence gGFP (F2-R2) found in higher concentrations *in planta*. Upon running the reactions on a denaturing gel we found that in the presence of both RppH and XRN-1 the larger F1-R2 is degraded to what appears to be completion (**Fig 5A**). Indeed the gGFP specific transcript was completely recalcitrant to degradation regardless of enzymes added (**Fig 5A**). Furthermore when reactions were ran on a non-denaturing agarose gel we found that XRN1 containing reactions migrated at a slower rate than the sample without XRN-1, possibly indicating XRN-1/gGFP binding, and also without gGFP degradation (**Fig 5B**). While this was a rather remarkable hypothesis-supporting result we still questioned if, in fact, sgRNAs transcribed *in vivo* displayed the same property. Therefore, cytosolic Cas9-IP samples (reported in **Fig 3B**) were treated with XRN-1 followed by RT-PCR to amplify a gGFP specific product. While there might have been a slight reduction in band intensity in the XRN-1 sample compared to the mock sample, there was certainly a substantial population of RNAs that remained resistant to the treatment (**Fig 5C**). Taken in total we believe this demonstrates XRN-1 resistance of the mature sgGFP transcripts, indicating a potential mechanism that native 5′ to 3′ exoribonucleases play a role in 5′ processing of progenitor-gGFP seen *in planta*.

**Figure 5.**
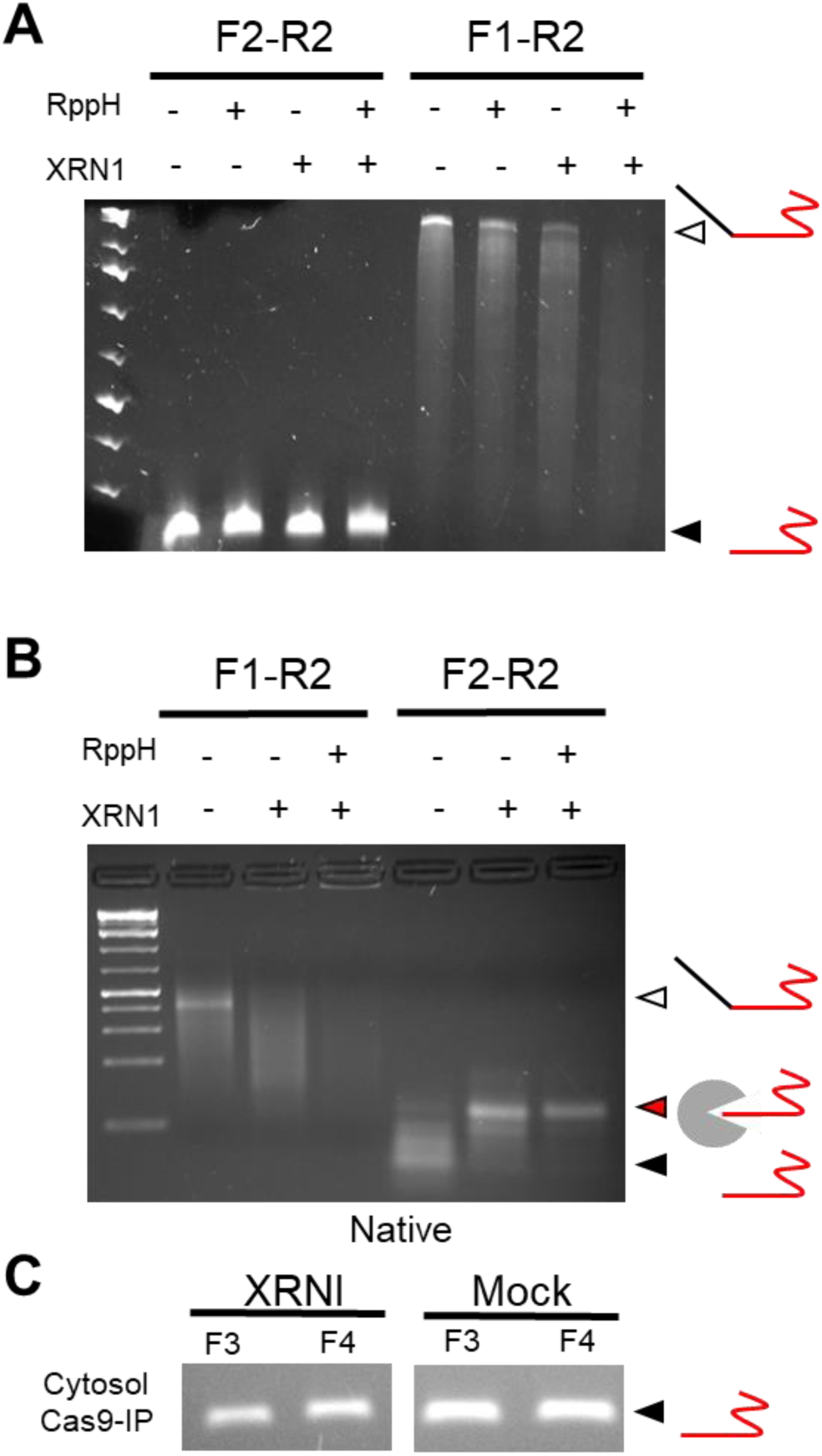
*In vitro* activity of XRN1 protein on GFP-gGFP and gGFP transcripts generated *in vitro* and *in planta.* **A)** *In vitro* assays using sgRNA transcripts containing 5′ overhangs (F1-R2 empty triangle) or without (F2-R2 black filled triangle) ran on a denaturing polyacrylamide gel. Transcripts were incubated in the presence of either no enzyme, the 5′ phosphatase RppH, 5′ to 3′ exonuclease XRN1 or both. **B)** *In vitro* assays using the experimental setup described in **A**. Reactions were ran on a native agarose gel to preserve transcript structure. Predicted transcript structure is demonstrated on the right side of the gel. Labeling remains the same as seen in **A**. However, the triangle with a red fill indicates undigested and possibly bound sgRNA transcripts by XRN1. **C)** RT-PCR from *in planta* generated sgRNA-Cas9 bound transcripts from the cytosolic fraction shown in figure **3B** subjected to XRNI or Mock enzyme treatment. F3-R2 and F4-R2 primers (**2B**) were used to amplify following treatments (total RNA and cytosolic RNA controls can be seen in **3B**).

## Discussion

While other studies have analyzed the 3’ processing of CRISPR RNAs (crRNAs) and trans-activating CRISPR RNAs (tracrRNAs) in both native and non-native CRISPR systems by RNAse III enzymes (Deltcheva et al. 2011; Karvelis et al. 2013), the mechanism of 5’ processing of crRNA/tracrRNAs or sgRNAs remains unknown (van der Oost et al. 2014). Even though it was highlighted that a secondary processing step focused on 5’ overhang removal must be taking place (van der Oost et al. 2014; Deltcheva et al. 2011) insight into the mechanism for sgRNA processing is largely unknown, and certainly no implication have been considered that it could be conserved amongst phylogenetic kingdoms. Furthermore, considerable amount of attention has been placed on nucleotide mismatches in the 20 nt complementary spacer sequence of sgRNAs and the associated loss of catalytic capabilities of the subsequent Cas9-sgRNA complexes on protospacer sequences (Jinek et al. 2012; Zheng et al. 2017). However, in depth studies of extraneous nucleotides 5′ to the sgRNA sequence and the biological effect on Cas9/sgRNA DNA cleavage events has not been reported to date.

In this study we systematically examined the effect of both 5′ and 3′ overhangs on Cas9- sgRNA complexes through a series of *in vitro* assays, which determined that specifically overhangs 5′ proximal to the sgRNA sequence inhibited the catalysis (DSB creation) of protospacer carrying DNA (the target for sgRNA). Following these results we hypothesized that in order for TRBO-G-3′gGFP to be catalytically active *in planta*, processing of the nucleotides 5′ proximal to the sgRNA sequence must occur. Furthermore, we demonstrated that transcripts produced from TRBO-G-3′gGFP resulted in an enrichment for 5′ processed sgRNA (5′ nucleotides not corresponding to the sgRNA removed) products bound by Cas9 in leaf tissue co-expressing both pHcoCas9 and TRBO-G-3′gGFP. Additional experimentation localized the preferred site of the Cas9 bound sgRNA processing events to the cytosol.

We next inquired if the sgRNA processing events were based on the cytosolic expression of TRBO subgenomic RNAs and/or if it was a viral-dependent RNA processing event. The utilization of U6 promoter driven expression of the protein ORF-sgRNA fusion transcript, GFP- gGFP, corroborates that cytosolic expression is optimal, however, possibly not strictly necessary for sgRNA transcript processing, and that these events are not unique to TRBO delivery of sgRNAs. In an attempt to obtain a better idea of what cellular pathway might be responsible for 5′ sgRNA processing we took an *in vitro* approach based on previous literature precedence which indicated a resistance of the native *S. pyogenes* tracrRNAs (analogous to the sgRNA “scaffold” sequence or the 3′ most 80 nucleotides) to 5′ to 3′ exonucleases (Deltcheva et al. 2011). We found that both *in vitro* and *in planta* transcribed gGFP RNA (or sgRNA of choice) demonstrated resistance to the 5′ to 3′ exonuclease XRN-1.

In **Figure 6** we suggest a model containing two not mutually exclusive parallel pathways for 5’ sgRNA processing for cytosolic transcribed (viral) RNAs in *N. benthamiana* based on results from experiments presented here (**Fig 6A**). Even though this model explains the nuclear generated transcripts processing events, we focused on the viral delivery for simplicity. Upon viral expression of transcripts that contain 5’ sequences that do not correspond to the sgRNA sequence, cytosolic localization is critical for optimal processing after or prior binding by Cas9. In order for sgRNA transcripts to be trimmed to the correct length, Cas9 binding might be necessary, as has been suggested previously (Mikami et al. 2017), or due to the sgRNA being inherently recalcitrant to exonuclease (**Fig 6B**). Specifically, on one hand Cas9 binding can be inferred to as important for proper processing (**Fig 2C** and **2D**), which agrees with previous structural analysis of the Cas9-sgRNA complex (Nishimasu et al. 2014; Anders et al. 2014) demonstrating that the 5′ most end of the sgRNA sequence is located within the active site of Cas9. In essence, the inclusion of the sgRNA sequence within the protein would protect the RNA from further degradation by host ribonucleases. This leads to one theory for our observed *in planta* catalytic activity in which Cas9 “shields” the sgRNA sequence from further degradation by exo- or endoribonucleases (right panel **Fig 6B-D**). However, we also provide evidence that sgRNAs are resistant to at least one class of nucleases, the 5′ to 3′ exonucleases (**Fig 5**). Due to the reliance of Cas9/sgRNA duplex catalytic activity on the 5′ sequence specificity of the sgRNA (**Fig 1C**), we find this result particularly relevant. While endoribonucleases might affect the formation of catalytically active Cas9-sgRNA complexes it seems rather unlikely that an endonuclease would have the specificity seen in **Figure 2F** or to produce catalytic events at the rate seen in **Figure 4C**.

**Figure 6.**
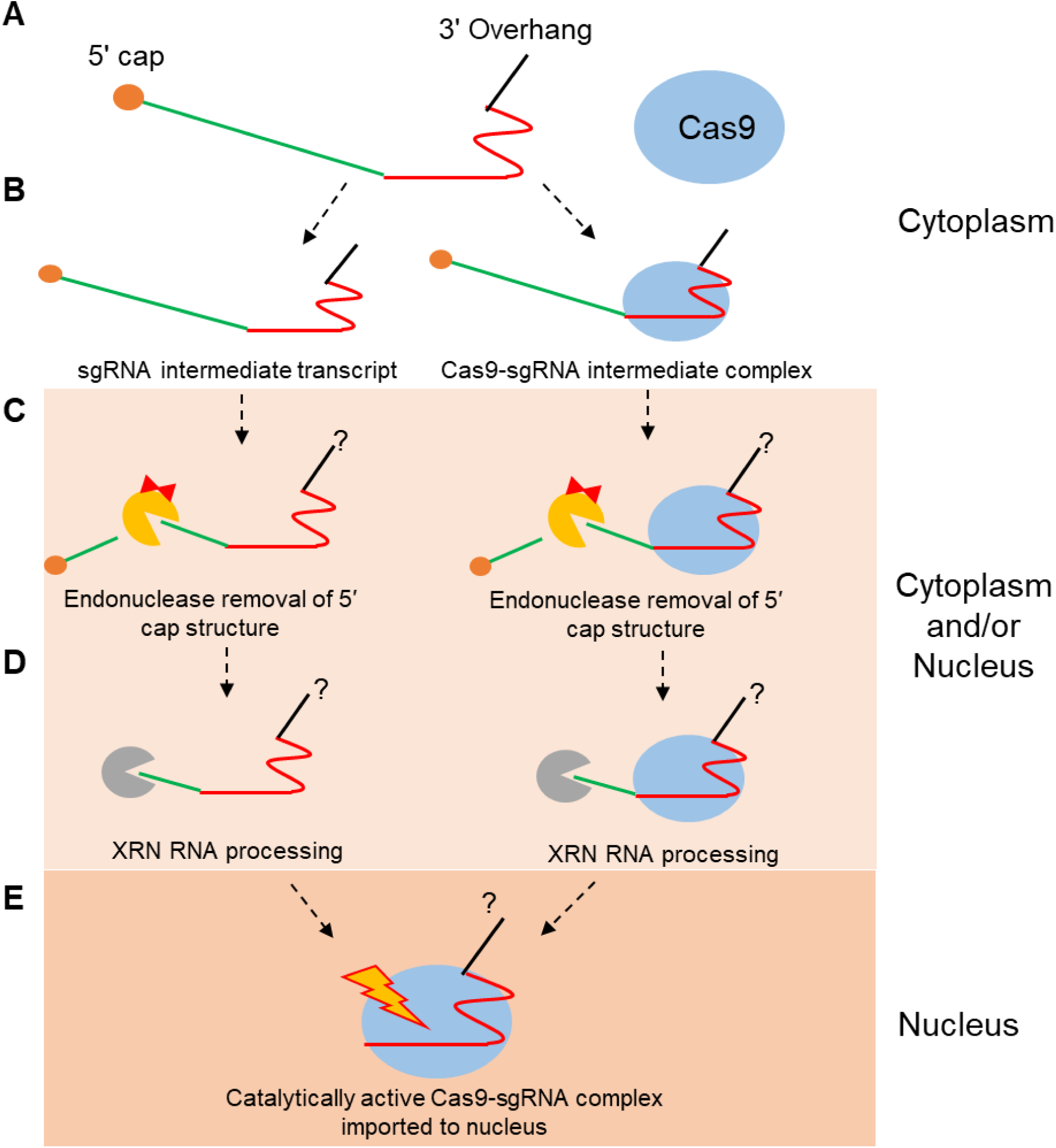
Contemporary model for sgRNA 5′ processing of GFP-gGFP in *N. benthamiana.* Transcription of the capped (orange sphere) transcript of GFP (green line) gGFP (red line) and the TMV 3′ UTR (back line) fusion transcript occurs in the cytoplasm of TRBO-GFP-gGFP infected cells (**A**). Following transcription the sgRNA containing transcript is either bound to Cas9 or remains unbound in the cytoplasm. Cytosolic endonucleases first remove the 5′ cap structure of the transcript (**B**) followed by either cytosolic or nuclear processing of the 5′ proximal nucleotide overhang through endogenous exonucleases (XRN1-like) occurs (**D**). Catalytically active Cas9- sgRNA complex is transported into the nucleus or remains in the nucleus as a catalytically active complex (**E**).

In either one of the above cases it seems that the most likely scenario for the processing events, at least using our transcriptional models (TRBO-G-3′gGFP and pHco-U6-GFP-gGFP), is an initial RNA cleavage event by an endoribonuclease which would provide the essential step of removing the 5′ mRNA cap and thus exposing the 5′ monophosphate group necessary for exoribonuclease activity (Jinek et al. 2011). The involvement of at least one endoribonuclease pathway, RNA silencing, was supported by preliminary results using the P19 RNAi suppressor in the experiments. This may not be terribly surprising considering the intimate relationship documented between RNAi and the 5′ to 3′ RNA degradation pathways, including XRN proteins involvement (Kurihara 2017). We believe that this model is further supported by our previous report (Cody et al. 2017) demonstrating the multiplexed delivery of two sgRNAs on a single viral transcript which lacked an autocatalytic (ribozyme) or ribonuclease specific sequences, such as the tRNA or Csy4 recognition sequence (Cermak et al. 2017b; Xie et al. 2015). The most reasonable interpretation of our previous success using multiplexed sgRNA delivery on a single cytosolic transcript, along with data presented in this manuscript, is that *N. benthamiana* first cleaves 3′ proximal to the first sgRNA, through an undetermined pathway, followed by XRN 5′ processing of the second sgRNA, yielding two biologically active sgRNAs. The importance of exoribonuclease degradation of sgRNAs is further supported by current structural knowledge of the Cas9/sgRNA complex, albeit circumstantial (Nishimasu et al. 2014; Anders et al. 2014). However, here we support a more important role for the XRN family and their processing of sgRNA. We believe that the recalcitrance of sgRNAs, and tracrRNAs (Deltcheva et al. 2011), is not merely coincidence, but instead an essential part of the native maturation system.

The translational impact of this report reaches beyond the fields of basic CRISPR biology or plant biology. It has been shown previously that there are native mechanisms for processing CRISPR arrays in non-native bacterial systems (Sapranauskas et al. 2011), human cells (Cong et al. 2013) as well as in plants (Mikami et al. 2017; Cody et al. 2017). However, these findings seem to have been overlooked in a multitude of studies that led to extravagant engineering of *in vivo* sgRNA delivery platforms; these have perhaps been developed based on the incorrect premise that, outside of the native *S. pyogenes* system, CRISPR-Cas9 delivery must be supplemented with specialized sgRNA delivery tools (Xie et al. 2015; Tsai et al. 2014; Gao and Zhao 2014; Cermak et al. 2017a). Perhaps one explanation for this development is the inherent focus on the 3′ processing of crRNAs by the RNase III enzyme in bacteria (Deltcheva et al. 2011; Sapranauskas et al. 2011) and human cells (Cong et al. 2013; Hsu et al. 2014) that appear to have a functional overlap among prokaryotes and eukaryotes. However, similar discussions about the secondary processing step of 5′ ends of crRNAs or sgRNAs *in vivo* are not as evident. Nevertheless, further evidence found in the native CRISPR type II-C system crRNA synthesis demonstrates the dispensable nature of the RNase III enzyme for creating catalytic Cas9- crRNA/tracrRNA complexes in *Neisseria meningitides* (Xu et al. 2013). The ability of *N. meningitides* and *Campylobacter jejuni* to produce crRNAs that do not appear to need 5′ processing through the intrinsic specificity of the 5 transcriptional start site of the crRNA promoter, we believe, further implicates the importance on the 5′ processing step for catalytic or interference activity of the complex (Dugar et al. 2013). RNase III cleaving of Cas9/crRNA/tracrRNA complexis from the native crRNA-spacer array can serve as a method for separation of crRNA/tracrRNA duplexes from a single transcript, but to be catalytically “activated” processing of the crRNA 5′ end still must occur. We speculate that this could occur through an analogous 5′ to 3′ exoribonuclease, such as RNase J1, which has shown a similar activity as demonstrated here, in *Bacillus subtilis* in regards to 16S ribosomal RNA processing (Mathy et al. 2007).

Perhaps the excitement of the potential uses of CRISPR systems has exceeded our knowledge of its basic biological processes. However, with this study the spotlight might shift to a realization of the ingenious engineering feat used by bacteria to harness established highly conserved native RNA degradation pathways for multiple cellular tasks.

## Materials and Methods

### Cloning and construct development

pHcoCas9 and TRBO-G-3′gGFP were constructed, explained, demonstrated to effectively cleave genomic DNA *in planta* and consistent indel detection through sequencing, in detail previously (Cody et al. 2017). The p-NLSCas9 plasmid was constructed using the human codon-optimized Cas9 nuclease (HcoCas9) (Addgene: 42230) (Cong et al. 2013) as a template. For this, the Cas9 encoding sequence without the NLS was amplified using a forward primer designed downstream of the NLS sequence and contained a *Bam*HI site as well as a start codon (ATG) and a reverse primer was designed upstream of the C terminal NLS sequence followed by a stop codon (TAA) and an *Xho*I site. The PCR amplicon was then cloned into a modified pRTL22 (Restrepo et al. 1990) sub-cloning vector, as previously. The 35S-Cas9-term cassette was then transferred into the binary destination plasmid pBINPLUS-sel using the *Hin*dIII site to create p-NLSCas9.

pHco-U6-gGFP and pHco-U6-GFP-gGFP were constructed using the pChimera subcloning vector Fauser et al. (2014) as a template for amplifying the U6 promoter and terminator for Gibson assembly into *Pac*I linearized pHcoCas9. TRBO-G-3′gGFP was used to amplify both gGFP for pHco-U6-gGFP and GFP-gGFP for pHco-U6-GFP-gGFP constructs.

### Cas9/sgRNA *in vitro* cleavage assays

TRBO-G-3′gGFP RNA templates containing either 5′ and 3′, 5′ or 3′, or non-overhang nucleotides flanking gGFP templates were synthesized using T7 RNA synthesis (New England BioLabs, Cambridge, MA). T7 RNAs were synthesized using 150 ng of each PCR template amplified from TRBO-G-3′gGFP. Forward primers T7-F1 and T7-F2 contained a T7 promoter followed by either the start sequence of GFP or gGFP, respectively. Reverse primers R1 and R2 corresponded to sequence in the TMV 3′ UTR or the 3′ most end of the sgRNA scaffolding sequence, respectively. RNA synthesis reactions were verified using 1% agarose gel electrophoresis stained with ethidium bromide and quantified using a NanoDrop (ThermoFisher scientific, Waltham, MA).

A PCR *mgfp5* fragment was amplified from untreated 16c genomic DNA followed by a cleanup step using DNA Clean & Concentrator −5 (Zymo Research) kit. Subsequently, 100 nM (final concentration) of purified Cas9 Nuclease was first incubated in Cas9 Nuclease reaction buffer (New England Biolabs, Cambridge, MA) and 30 nM (final concentration) with each of the T7 synthesized gGFP containing transcripts, or for assays using varying concentrations of gGFP template with the corresponding final nM concentrations indicated, and added to each reaction and incubated at room temperature for 5 minutes. Then, 3 nM (final concentration) of purified *mgfp5* PCR template was added to each reaction and incubated for 60 minutes at 37°C. Reactions were visualized using 1.5 % agarose gel electrophoresis stained with ethidium bromide.

### Agroinfiltrations

*Agrobacterium tumefaciens* strain GV3101 (pMP90RK) was used for binary plasmid infiltrations in *N. benthamiana*, as previously described (Odokonyero et al. 2015). In brief, GV3101 cultures were grown overnight (16-20 hrs) under 250 rpm shaking at 28 °C in LB media supplemented with 50 mg/L Kanamycin. Cells were pelleted through centrifugation and suspened in infiltration buffer (10 mM MgCl_2_, 10 mM MES pH 5.7, and 200 µM acetosyringone). TRBO-based cultures were resuspended to a final infiltration concentration of OD_600_ 0.4 and Cas9 expression vectors (pHcoCas9 and p-NLSCas9) at OD_600_ 0.5. 16c plants that were four week old were used for *Agrobacterium* infiltrations of the abaxial side of the leaf. Plants were returned to normal growth conditions.

### DNA and indel assays

Single plant DNA samples for indel assays were carried out using leaf tissue from three infiltrated leaves, totaling 100-150 mg of tissue, to serve as pooled biological replicates and to avoid tissue-dependent effects. DNA extractions were then carried out using *Quick* DNA Miniprep kit (Zymo Research). A total of 100 ng of genomic DNA was used for PCR amplification of *mgfp5* gene from 16c plants. Due to the previous consistent results of amplicon sequencing and *Bsg*I assay results we used this assay as our indel detection method of choice. Amplicons were cleaned using DNA Clean & Concentrator −5 (Zymo Research) kit and 250-400 ng DNA was used for *Bsg*I digestions which were incubated at 37°C overnight. *Bsg*I restriction enzyme resistance assays were then visualized using 1.2% agarose gel electrophoresis stained with ethidium bromide. Image files (.tif) were uploaded in the image analysis software ImageJ (NIH) and band intensities were measured using gel peak analysis.

### Cas9-gRNA immunoprecipitation assays

At 3 dpi *N. benthamiana* tissue was ground in liquid nitrogen and resuspended in RIPA buffer (50 mM Tris-HCl pH 8.0, 2 mM EDTA, 1% Triton X-100, 0.1% SDS, 0.5% Na-deoxycholate, and 150 mM NaCl) at a ratio of 1g of tissue to 3 ml of buffer. Tissue was centrifuged at 10,000 g for 20 min., and supernatant filtered through Miracloth (EMD Millipore) pre-soaked in RIPA buffer. Then, 4 ml of lysate was incubated by end over end agitation in 4°C for 4 hrs with 1:400 anti-Cas9 antibody (Biolegend). Following, 200 µl of protein G agarose slurry (Thermo Scientific) was added and the mixture was incubated for an additional hour at 4°C. Cas9-protein G beads were collected through centrifugation at 2,500 g for 3 min. Supernatant was removed and the agarose slurry was washed 5 times with 500 µl of RIPA buffer. Following wash steps some of the resuspended slurry was used for western blot analysis to detect proper Cas9-immunoprecipitation and the rest was used for RNA extractions.

### Nuclei isolation

Leaf tissue was isolated at 3 dpi and ground in liquid nitrogen and resupended at a ratio of 1g to 5 ml of nuclei isolation buffer (0.25 M, sucrose, 15mM PIPES pH 6.8, 5 mM MgCl_2_, 60 mM KCl, 15 mM NaCl, 1 mM CaCl_2_, 0.9% Triton X-100). The ground tissue and buffer solution was then incubated on ice for 30 min. Following incubation, the sample was centrifuged at 10,000g for 20 min at 4°C. The supernatant (cytosol fraction) was used directly for assays while the pellet (nuclei fraction) was washed and re-isolated with 500 µl of nuclei isolation buffer 3 times. Clean nuclei pellets were resuspended in 100 µl of RIPA buffer and used for downstream assays. Successful subcellular isolation was demonstrated using Coomassie staining as previously described (Desvoyes et al. 2002).

### RNA extractions and (RT)-PCR

*N. benthamiana* RNA extractions were performed using the Direct-zol RNA Miniprep kit (Zymo Research) following the instructions from the manufacturer. cDNA was then synthesized with equal concentrations of total RNA or equal volumes of solution (RIP) using the M-MLV Reverse Transcriptase (Invitrogen) and gene specific primers. RT-PCR was carried out using Q5 High Fidelity Polymerase (New England Biolabs, Cambridge, MA).

### XRN1 degradation assays

*In vitro* XRN1 degradation assays were performed on F1-R2 and F2-R2 transcripts which were synthesized using T7 polymerase, as stated above. 300 ngs of each transcripts was then incubated with 0.5 units of XRN-1 both with and without 2.5 units of RppH using the manufactures supplied buffer for 1 hour at 37° (New England Biolabs, Cambridge, MA). Reactions were ran in either denaturing (4% Urea PAGE) or native conditions (agarose). Similarly, *in planta* synthesized transcripts from cytosolic Cas9-RIP assays were treated with or without (mock) 1 unit of XRN-1 and 20 units of RNAse inhibitor Murine and incubated at the above specified conditions. Reactions were then subjected to an RT and a PCR step to assay for the presence of sgRNA specific transcripts.

## Acknowledgements

We thank April DeMell, Kelvin Chiong and Maria Mendoza for input and assistance throughout the project, and K.-B. G. Scholthof, K. K. Mandadi and T. Erik Mirkov for their helpful critical comments and insightful discussions throughout the study or during manuscript preparation.

